# Nup358 regulates remodelling of ER-mitochondrial contact sites and autophagy

**DOI:** 10.1101/2021.10.01.462723

**Authors:** Misha Kalarikkal, Rimpi Saikia, Pallavi Varshney, Prathamesh Dhamale, Amitabha Majumdar, Jomon Joseph

**Affiliations:** National Centre for Cell Science, S.P. Pune University Campus, Pune - 411007, India

## Abstract

The contact sites between ER and mitochondria regulate several cellular processes including inter-organelle lipid transport, calcium homeostasis and autophagy. However, the mechanisms that regulate the dynamics and functions of these contact sites remain unresolved. We show that annulate lamellae (AL), a relatively unexplored subcellular structure representing subdomains of ER enriched with a subset of nucleoporins, are present at ER-mitochondria contact sites (ERMCS). Depletion of one of the AL-resident nucleoporins, Nup358, results in increased contacts between ER and mitochondria. Mechanistically, Nup358 modulates ERMCS dynamics by restricting mTORC2/Akt signalling. Our results suggest that growth factor-mediated remodelling of ERMCS depends on a reciprocal binding of Nup358 and mTOR to the ERMCS tethering complex consisting of VAPB and PTPIP51. Furthermore, Nup358 also interacts with IP3R, an ERMCS-enriched Ca^2+^ channel, and controls Ca^2+^ release from the ER. Consequently, depletion of Nup358 leads to elevated cytoplasmic Ca^2+^ and autophagy via activation of Ca^2+^/CaMKK2/AMPK axis. Our study thus uncovers a novel role for AL, particularly for Nup358, in regulating mTORC2-mediated ERMCS remodelling and Ca^2+^-directed autophagy, possibly via independent mechanisms.

## INTRODUCTION

Contrary to the general perception that organelles in the cell exist and function independently, recent developments highlight that multiple organelles make physical and dynamic contacts with each other to coordinate the many functions they perform (Henne, 2016; Prinz et al., 2020; Scorrano et al., 2019; Wu et al., 2018). Endoplasmic reticulum (ER), an organelle characterized by its membranous network that extends throughout the cytoplasm, interacts with other organelles through specific contact sites (Cohen et al., 2018; López-Crisosto et al., 2015; Petkovic et al., 2021; Phillips and Voeltz, 2016). ER-mitochondria contact sites (ERMCS), also known as membrane-associated mitochondria (MAM), are maintained through physical interactions between proteins present on both the organelles (Marchi et al., 2014; Raturi and Simmen, 2013; Rowland and Voeltz, 2012; Wu et al., 2018)(Filadi et al., 2017). Some of the ERMCS tethering complexes include VAPB-PTPIP51 (De vos et al., 2012), Mfn2-Mfn1/2 (De Brito and Scorrano, 2008), IP3R-GRP75-VDAC (Szabadkai et al., 2006) and BAP31-Fis1 (Iwasawa et al., 2011). Growth factor-stimulated mTORC2 pathway is shown to stabilize ERMCS through Akt-mediated phosphorylation of ERMCS proteins such as IP3R3 and PACS2 (Betz et al., 2013).

ERMCS regulate inter-organelle Ca^2+^ transfer, lipid transfer, energy metabolism, inflammation, apoptosis, autophagy and several other ER/mitochondrial functions (Barazzuol et al., 2021; Csordás et al., 2018; Madreiter-Sokolowski et al., 2019; Perrone et al., 2020; Vance, 2020). The ER-resident Ca^2+^ channel IP3R is enriched at the ERMCS where it mediates Ca^2+^ transfer from the ER to the mitochondria to regulate energy metabolism and apoptosis (Csordás et al., 2018; Grimm, 2012). Consistent with the range of processes these contact sites regulate, dysfunctional ERMCS are implicated in many disorders including diabetes (Rieusset, 2018), cancers (Doghman-Bouguerra and Lalli, 2019; Peruzzo et al., 2020; Simoes et al., 2020) and neurodegeneration (Paillusson et al., 2016; Petkovic et al., 2021). Although many cellular functions are dependent on ERMCS, the molecular interplay regulating the dynamic interaction between ER and mitochondria remains unclear.

The ER is involved in a multitude of physiological functions, which are performed by specialized subdomains of the organelle (Cohen et al., 2018). Annulate lamellae (AL), an underexplored subdomain of the ER, are stacked membranes containing pore-like assemblies that structurally resemble the nuclear pore complexes (NPCs) present on the nuclear envelope (NE) (Kessel, 1992). Many functions for AL have been proposed based on electron microscopic observations, which include roles in infection, cancers, gene expression, mRNA regulation and development (Kessel, 1992). AL are remodelled extensively during embryonic development, infection by intracellular pathogens including SARS-CoV-2 and in cancers (Eymieux et al., 2021; Kessel, 1992). Recently, involvement of AL in functional assembly of NPCs has been reported (Hampoelz et al., 2016). AL possess only a subset of nucleoporins as compared to the NPCs (Cordes et al., 1996; Sahoo et al., 2017), indicating a compositional, structural and functional difference between AL pore complexes and NPCs. A role for the AL-resident nucleoporin Nup358 in microRNA-mediated translational suppression of mRNAs has been documented (Sahoo et al., 2017).

Since ER makes contacts with mitochondria through specialized subdomains at the ERMCS, and AL represent another subdomain of the ER, we explored if these two subdomains functionally interact with each other. We found that Nup358-positive AL were often present at ERMCS. Interestingly, Nup358 negatively regulates ERMCS integrity through suppression of mTORC2/Akt activation and inhibits autophagy by restricting IP3R-mediated Ca^2+^ release into the cytoplasm.

## RESULTS

### Annulate lamellae are present at ER-mitochondrial contact sites

As AL define subdomains of ER, and the ER makes extensive contacts with multiple organelles including mitochondria, we tested the hypothesis that AL is present at ERMCS. Microscopic images of cells co-stained for Nup38 as an AL marker (Sahoo et al., 2017), PDIA3 as an ER marker and MitoTracker for mitochondria revealed that AL are often present at ERMCS (**Fig. 1a**). Consistent with this, stimulated emission depletion microscopy (STED) of AL (Nup358) and mitochondria (Tom-20) also showed that many of the AL structures were present adjacent to mitochondria (**Fig. 1b**). Fractionation of organelles from HeLa cells confirmed that a pool of Nup358, along with other nucleoporins (Nup62 and Nup88), was present in the mitochondria associated membrane fraction, which represents ERMCS (Montesinos and Area-Gomez, 2020) (**Fig. 1c**). Disruption of ERMCS by depletion of specific proteins such as VAPB-PTPIP51, Mitofusin 2 and IP3R3, which were previously shown to be required for ERMCS integrity led to disappearance of AL, as judged by absence of Nup358 and other AL-associated nucleoporins **(Supplementary Fig. 1a)**. However, localization of these nucleoporins to the nuclear membrane remained unaffected under the above condition. Since Nup358 is an authentic AL marker (Sahoo et al., 2017), we conclude that AL reside at ERMCS and their assembly may depend on ERMCS integrity.

**Fig. 1.**
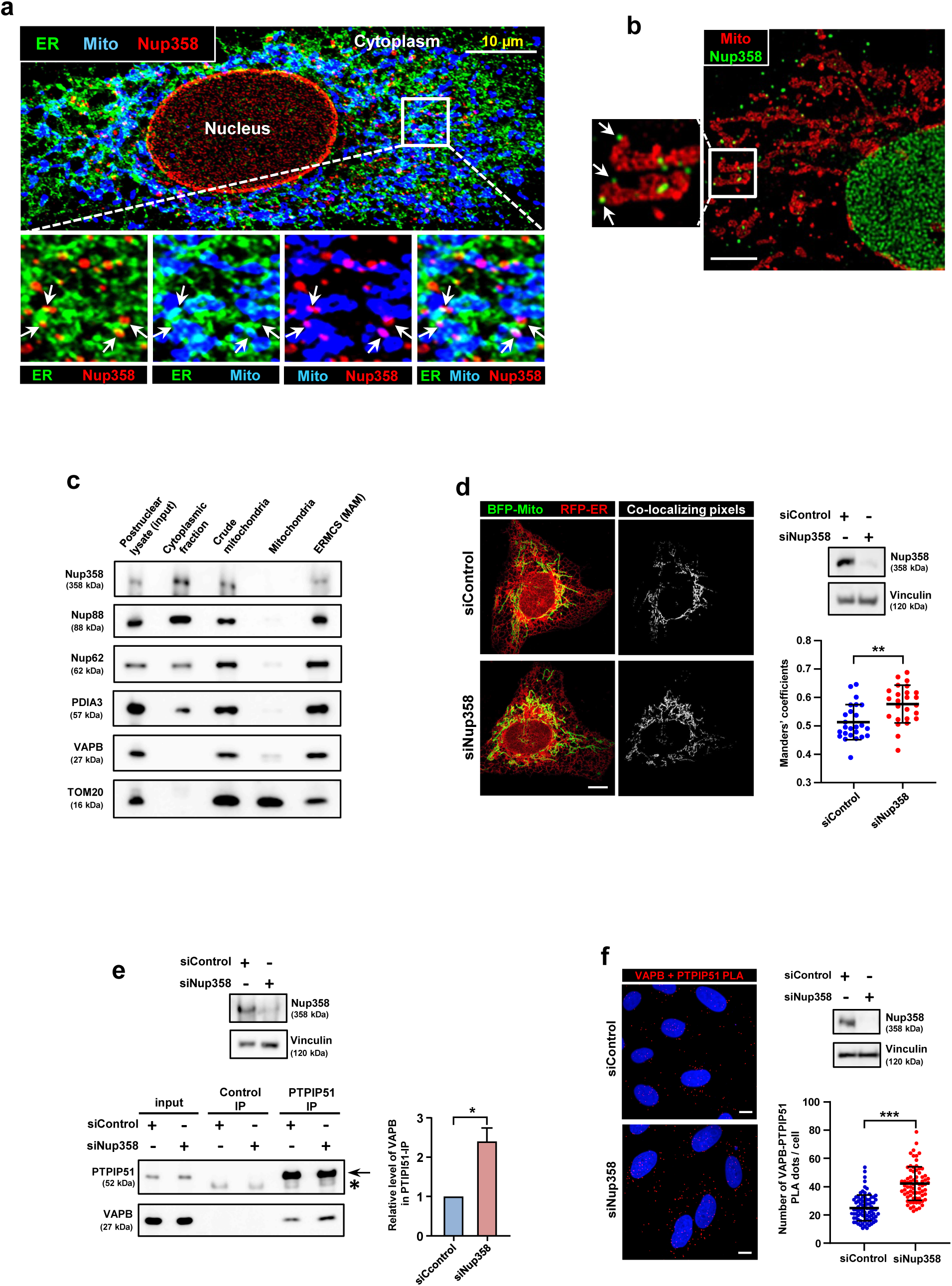
AL-resident Nup358 is present at the ERMCS and regulates their dynamics. **a**, Nup358-positive AL are present at ERMCS. Confocal microscopic image displaying the relative localization of indicated proteins; Nup358 for AL (red), PDIA3 for ER (green) and MitoTracker for mitochondria (blue) in U2OS cells. Arrows show Nup358-positive AL. Scale bar, 10 μm. **b**, Stimulated emission depletion super-resolution microscopy of Huh-7 cell immunostained with AL marker (Nup358, green) and mitochondria marker (Tom-20, red). Scale, bar 5 μm. **c**, Nup358, along with Nup88 and Nup62, is present in ERMCS fraction. HeLa cells were processed to obtain ERMCS (MAM) fractions, which along with other fractions were analysed for specific proteins by western blotting. **d**, Nup358 depletion leads to increased contacts between ER and mitochondria. U2OS cells were initially transfected with control (siControl) or Nup358 (siNup358) specific siRNA and later co-transfected with RFP-ER (green) and BFP-Mito (pseudo-coloured in green) constructs for labelling ER and mitochondria, respectively. Left: the co-localizing pixels are shown in grey. Scale bar, 10 μm. Right top: the extent of Nup358 depletion was evaluated by western blotting with specific Nup358 antibody. Vinculin was used as loading control. Right bottom: Analysis of individual Meanders coefficient values of RFP-ER with BFP-Mito from siControl and siNup358 treated HeLa cells (*n* = 25 cells from 3 independent experiments). Data are mean±SD, Mann-Whitney test. ** *P*≤0.01. **e**, In Nup358-deficient cells association of VAPB with PTPIP51 is enhanced. Extent of VAPB association with PTPIP51 was analysed by co-immunoprecipitation (co-IP) assay. Top: extent of Nup358 depletion assessed by western blotting. Bottom left: Co-IP of endogenous VAPB with endogenous PTPIPI51 from HeLa cells treated with siControl or siNup358. Bottom right: quantitation of the amount of VAPB associated with PTPIP51 (*n* = 3 independent experiments). Data are mean±SEM, unpaired Student’s *t* test. \**P*≤0.05 **f**, Enhanced interaction between VAPB and PTPIP51 in the absence of Nup358. In situ proximity ligation assay (PLA) was performed for assessing the interaction between VAPB and PTIPI51 using specific antibodies. Left: representative images showing PLA puncta (red) in HeLa cells treated with siControl or siNup358. Scale bar, 10 μm. Right top: extent of Nup358 depletion as analysed by western blotting, along with vinculin as loading control. Right bottom: quantitation of number of PLA puncta per cell from siControl and siNup358 HeLa cells (*n* = 82 cells for siControl and 73 cells for siNup358 from 3 independent experiments). Data are mean±SD, Mann-Whitney test. *** *P*≤0.001.

### Nup358 depletion increases the contacts between ER and mitochondria

We tested if the AL component Nup358 affects the contacts between the ER and the mitochondria by monitoring the ERMCS in Nup358 depleted cells in multiple ways. Knockdown of Nup358 in U2OS, an osteosarcoma cell line, led to enhanced contacts between ER and mitochondria as assessed by increased co-localization between ER and mitochondria by fluorescence microscopy (**Fig. 1d**). Moreover siRNA-mediated depletion of Nup358 from HeLa cells led to increased interaction between components of the ERMCS tethering complex, VAPB (ER) and PTPIP51 (mitochondria), as monitored by co-immunoprecipitation assays (co-IP) (**Fig. 1e**). The increase in ERMCS in the absence of Nup358 was further verified by proximity ligation assay (PLA) using antibodies against VAPB and PTPIP51, in HeLa cells (**Fig. 1f, Supplementary Fig. 2a**). However, the absence of Nup358 did not affect the intactness of AL (**Supplementary Fig. 1b**).

Previous studies have shown a role for Nup358 in the miRNA pathway (Sahoo et al., 2017; Shen et al., 2021). Disruption of the miRNA pathway via depletion of GW182 (Pfaff and Meister, 2013) did not affect ERMCS dynamics (**Supplementary Fig. 2b**). Collectively, these results show that Nup358 negatively regulates ERMCS stability independent of its function in miRNA pathway.

### Nup358 depletion-mediated increase in ERMCS is dependent on mTORC2/Akt signalling

Growth factor stimulation results in recruitment of mTORC2 to ERMCS, which via Akt-mediated phosphorylation of proteins such as PACS2 stabilizes the ERMCS (Betz et al., 2013). We examined the possibility that mTORC2/Akt signalling enhances the contacts between ER and mitochondria in Nup358-deficient cells. Interestingly, siRNA-mediated depletion of Nup358 from HeLa cells led to the activation of mTORC2, determined by the phosphorylation of its downstream kinase Akt at S473 (**Fig. 2a**). Activation of mTORC2 was confirmed in Nup358 knockout (KO) HeLa cells (**Fig. 2b, Supplementary Fig. 3**). Co-depletion of Rictor, a key subunit of mTORC2, reversed the hyper-phosphorylation of Akt resulting from growth factor [insulin and epidermal growth factor (EGF) signalling in the absence of Nup358 (**Fig 2c**). Knockdown of Nup358 also activated mTORC1, indicated by the phosphorylation of Akt at T308, S6K at T389 and S6 at S235/S236, which could be rescued by co-depletion of Rictor (**Fig. 2d**), suggesting that the hyperactivation of mTORC1 in Nup358-deficient cells depends on mTORC2/Akt activity (Szwed et al., 2021). Further, we tested if elevated mTOCR2/Akt activity increased the ERMCS in the absence of Nup358. Co-depletion of Rictor reversed the increase in ERMCS observed in Nup358 knockdown cells (**Fig. 2e**). Collectively, these results suggest that Nup358 negatively regulates ERMCS by suppressing mTORC2/Akt activation.

**Fig. 2.**
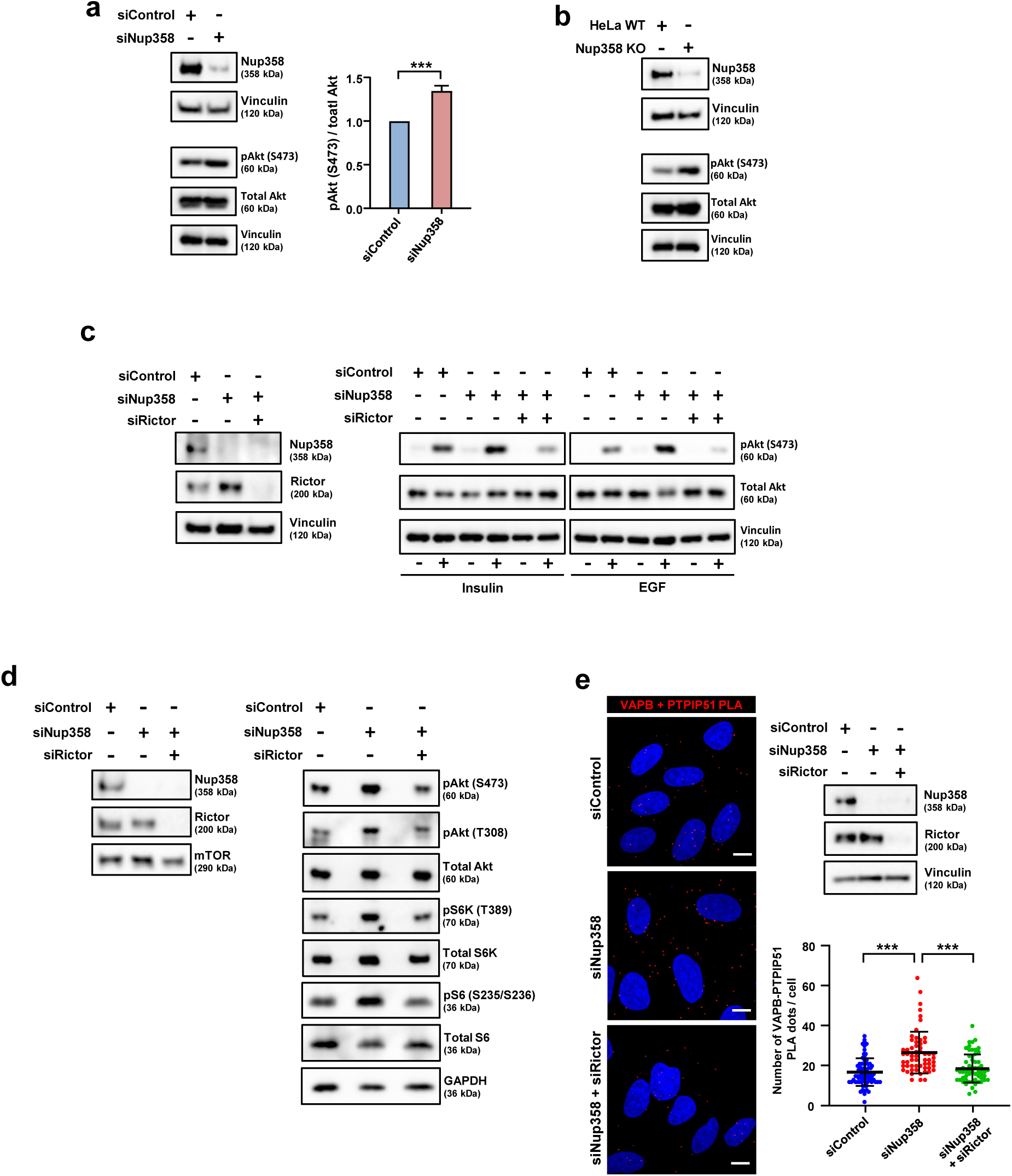
Increased ERMCS upon Nup358 loss is dependent on mTORC2/Akt signalling. **a**, Nup358 depletion leads to elevated mTORC2/Akt activation. Control (siControl) and Nup358 (siNup358) specific siRNA transfected HeLa cells were analysed for the extent of mTORC2/Akt activation by western blotting using antibodies as described (left). Vinculin was used as loading control. Right: quantitative representation of the relative level of pAkt (S473) as compared to total Akt under the above mentioned experimental condition (*n* = 3 independent experiments). Data are mean±SEM, unpaired Student’s *t* test. \**P*≤0.05. **b**, The hyperactivation of mTORC2/Akt signalling occurs in Nup358 knockout (KO) HeLa cells as compared to wild-type (WT) cells. **c**, Nup358 restricts mTORC2-mediated hyper-phosphorylation of Akt at S473 upon growth factor signalling. HeLa cells, transfected with indicated siRNAs, were serum starved for 3 h. Then the cells treated with (+) or without (-) Insulin (1 nM for 20 min) or epidermal growth factor (EGF, 20 ng/ml for 10 min were analysed for mTORC2/Akt activation by western blotting. Left: shows extent of depletion of specific proteins, with vinculin as a loading control. Right: shows the extent of Akt phosphorylation at S473 under different conditions as described. Vinculin was used as loading control. **d**, Nup358 depletion leads to mTORC1 activation, which was rescued by co-depletion of Rictor. HeLa cells treated with specific siRNAs were analysed for the extent of protein depletion using western blotting (left) and the extent of mTORC1 activation (right), as assessed by phosphorylation of indicated mTORC1 targets using western blotting. **e**, Increased contacts between ER and mitochondria upon Nup358 depletion depends on mTORC2 activity. HeLa cells were depleted of indicated proteins using specific siRNAs and analysed for the extent of ER-mitochondria contacts using *in situ* PLA with VAPB and PTPIP51 antibodies (left). Scale bar, 10 μm. The extent of protein depletion (right top) and quantitation of PLA dots per cell (right bottom) are shown (*n* = 72 cells for siControl; 59 cells for siNup358; 58 cells for siNup358 + siRictor from 3 independent experiments). Data are mean±SD, Mann-Whitney test. *** *P*≤0.001.

### ERMCS tethering complex VAPB-PTPIP51 regulates growth factor-stimulated mTORC2/Akt activation

The mTORC2 complex and Akt are recruited to ERMCS in response to growth factor signalling (Betz et al., 2013), and intact ERMCS are required for their activation (Tubbs et al., 2014). To check if specific proteins at the ERMCS are involved in the recruitment and/or activation of mTORC2/Akt, different proteins that localize to and regulate ERMCS such as VAPB/PTPIP51, PACS2, IP3R3 and Mfn2 were depleted in HeLa cells, and the activation status of mTORC2/Akt was monitored. Interestingly, loss of VAPB and PTPIP51, but not others, significantly reduced insulin-stimulated mTORC2/Akt activation (**Fig. 3a**). These results suggest that the VAPB-PTIP51 complex plays a specific role in growth factor-stimulated activation of mTORC2/Akt signalling.

**Fig. 3.**
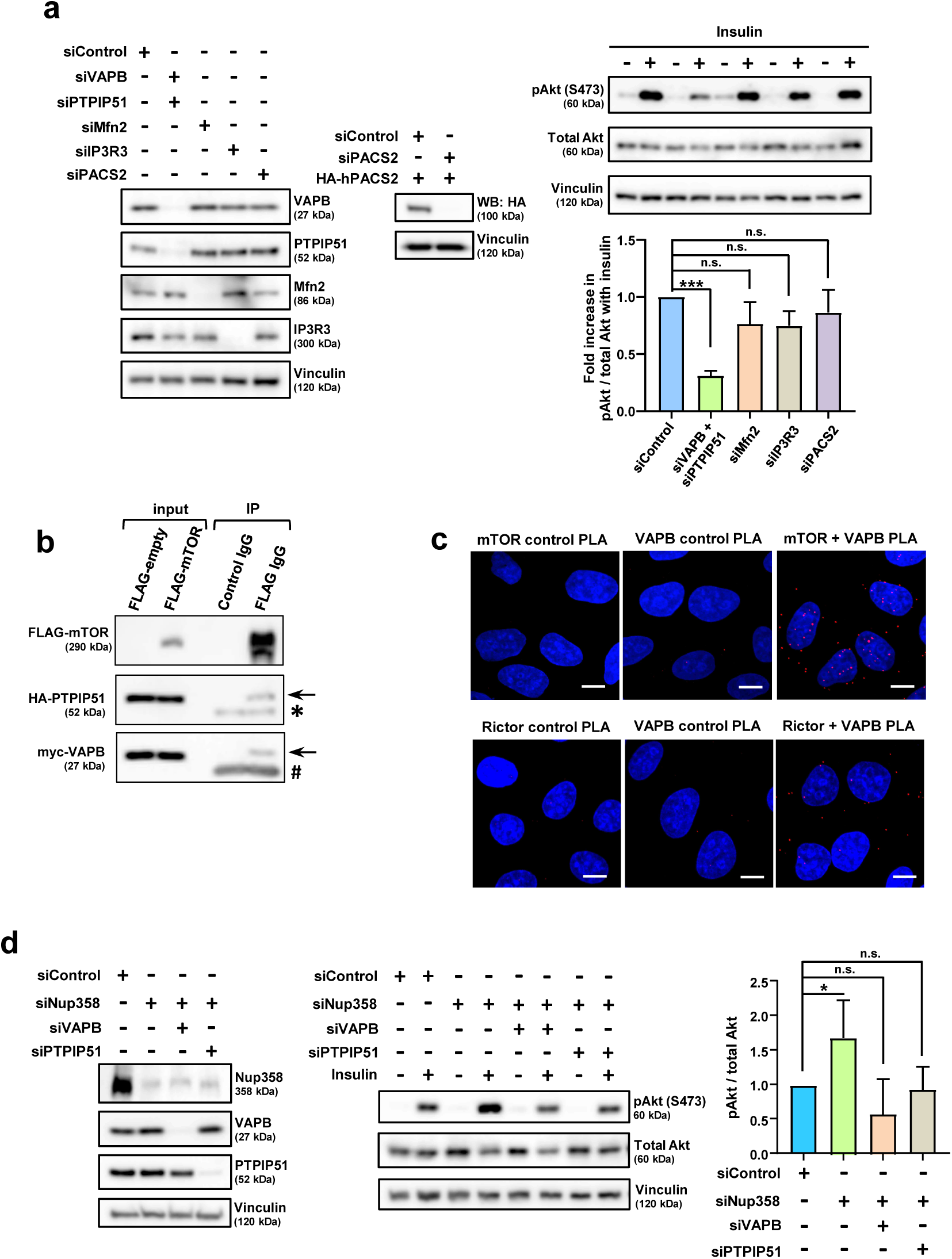
Hyperactivation of mTORC2/Akt in Nup358-deficient cells depends on VAPB-PTPIP51 tethering complex. **a**, Depletion of VAPB/PTPIP51 interferes with growth factor stimulated activation of mTORC2/Akt signalling. HeLa cells were depleted of different proteins as mentioned. The extent of depletion was verified by western blotting using specific antibodies (left). The ability of PACS2 siRNA to deplete the human PACS2 was confirmed by co-transfecting HeLa cells with control (sControl) or PACS2 (siPACS2) siRNA along with HA-human PACS2 construct. The expression of HA-PACS2 was monitored by western blotting (WB) with HA-specific antibody (middle). Vinculin was used as loading control. Right top: HeLa cells depleted of indicated proteins were serum starved for 3 h, and later treated with (+) or without (-) Insulin (1 nM) for 20 min. The cells were then analysed for the specific proteins by western blotting. Right bottom: quantitative analysis of the relative phosphorylation of Akt at S473 under the described conditions (*n* = 3 independent experiments). Data are mean±SEM, unpaired Student’s *t* test. n.s. not significant, *** *P*≤0.001. **b**, mTOR interacts with VAPB and PTPIP51. HEK293T cells expressing FLAG-control (Empty vector) or FLAG-mTOR along with HA-PTPIP51 and myc-VAPB were subjected to immunoprecipitation (IP) using FLAG-specific antibodies. Presence of HA-PTPIP51 and myc-VAPB in the IP samples was examined by western analysis. Arrow indicates HA-PTPIP51 or myc-VAPB as shown; * indicates IgG heavy chain and # indicates IgG light chain cross-reaction. **c**, Interaction between endogenous VAPB with mTOR (top) and Rictor (bottom) was confirmed by *in situ* PLA in HeLa cells. An increase in the number of PLA dots (red) is observed with mTOR or Rictor in combination with VAPB antibodies, as compared to single antibody controls (mTOR, Rictor or VAPB alone). Scale bar, 10 μm. **d**, Hyperactivation of mTORC2/Akt signalling upon Nup358 loss depends on the VAPB-PTPIP51 tethering complex. HeLa cells were depleted of indicated proteins and serum starved for 3 h. The cells were then treated with (+) or without (-) Insulin (1 nM) for 20 min. Cells were analysed for insulin-induced mTORC2/Akt activation. Left: Western analysis to monitor the extent of protein depletion by siRNAs. Middle: Western blot showing phosphorylation of Akt at Ser473 in the described conditions. Right: Quantitative data depicting the change in relative amount of pAkt (S473) upon insulin addition under indicated conditions (*n* = 3 independent experiments). Data are mean±SEM, unpaired Student’s *t*-test. n.s. not significant, \**P*≤0.05.

To test if the mTORC2 complex physically interacts with the VAPB-PTPIP51 complex, co-IP assay was performed. Indeed, myc-VAPB and HA-PTPIP51 co-immunoprecipitated with FLAG-mTOR (**Fig. 3b**). Moreover, specific endogenous interaction of mTOR and Rictor with VAPB was confirmed by PLA (**Fig. 3c**). Collectively, the data indicates that mTORC2 and VAPB-PTPIP51 complex physically associate with each other.

Based on our observation that the ER-mitochondria tethering complex VAPB-PTPIP51 is involved in growth factor induced mTORC2/Akt activation, we tested if the enhanced mTORC2 activity in the absence of Nup358 was mediated by VAPB-PTPIP51. Interestingly, depletion of VAPB or PTPIP51 rescued the hyperactivation of mTORC2/Akt signalling resulting from Nup358 knockdown (**Fig. 3d**), indicating that Nup358 negatively regulates mTORC2/Akt in a VAPB-PTPIP51 dependent manner. Additionally, the observation that depletion of either VAPB or PTPIP51 interfered with the mTORC2 hyperactivation in Nup358-deficient cells suggests that assembly of the VAPB-PTIPI51complex might be essential for mTORC2/Akt activation.

### Binding of Nup358 or mTOR to VAPB-PTPIP51 complex may determine the stability of ERMCS

Our data suggests that the ERMCS tethering complex VAPB-PTIPI51 mediates growth factor induced mTORC2/Akt activation and thereby strengthening of ER and mitochondrial contacts. Additionally, Nup358 negatively regulates ERMCS by suppressing mTORC2/Akt activation (**Fig. 2e**). Nup358 also interacts with VAPB and PTPIP51, as determined by co-IP and PLA (**Fig. 4a, b**).

**Fig. 4.**
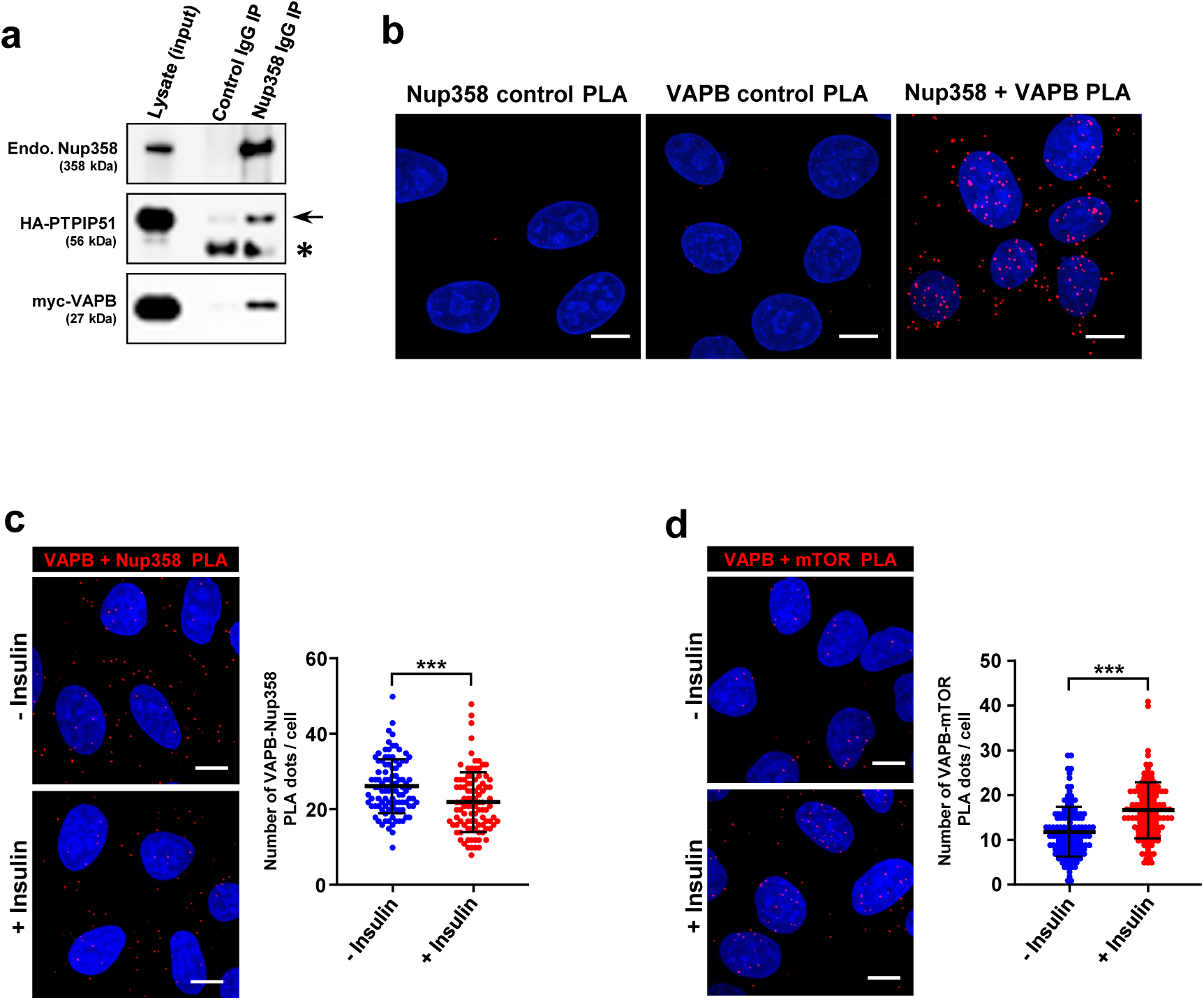
Reciprocal binding of Nup358 and mTOR to VAPB is modulated by growth factor signalling. **a**, Nup358 interacts with VAPB and PTPIP51. HEK293T cells expressing HA-PTPIP51 and myc-VAPB were subjected to endogenous Nup358 IP using specific antibodies. Presence of HA-PTPIP51 and myc-VAPB was examined by western analysis of the IP samples. Arrow indicates HA-PTPIP51 and * indicates IgG heavy chain cross-reaction. **b**, Interaction between endogenous Nup358 and VAPB was confirmed by *in situ* PLA using specific antibodies in HeLa cells. Nup358 or VAPB antibody alone was used as control. Scale bar, 10 μm. **c, d**, HeLa cells were serum starved for 3 h. Cells were later treated with (+) or without (-) Insulin (10 nM) for 20 min. Extent of Nup358 and VAPB interaction (**c**) or mTOR and VAPB interaction (**d**) was assessed by PLA under the described conditions. Left (**c, d**): representative microscopic images showing the PLA dots (red) with (+) or without (-) insulin treatment. Scale bar, 10 μm. Right (**c, d**): quantitative data depicting number of PLA dots per cell under indicated scenario. **c**, Number of Nup358-VAPB PLA dots per cell (*n* = 98 cells from 3 independent experiments) **d**, Number of mTOR-VAPB PLA dots per cell (*n* = 131 cells for Insulin untreated and 125 cells for Insulin treated from 3 independent experiments). Data are mean±SD, Mann-Whitney test. *** *P*≤0.001. Scale bar, 10 μm.

Considering that the VAPB-PTPIP51 complex associates with both Nup358 (**Fig. 4a, b**) and mTORC2 (**Fig. 3b, c**), we hypothesized that perhaps Nup358 when present at the ERMCS denies mTORC2 access to the VAPB-PTPIP51 complex, thereby limiting mTORC2 activity. Thus in the absence of growth factor signalling, Nup358 would inhibit the interaction between mTORC2 complex and VAPB-PTPIP51. On the contrary, growth factor signalling relieves the Nup358-mediated inhibition, thereby enhancing the interaction between VAPB-PTPIP51 and mTORC2, subsequently resulting in mTORC2 activation. To test this possibility, the kinetics of complex formation of VAPB with Nup358 or mTOR was monitored by PLA during insulin signalling. Consistent with the hypothesis, as compared to control conditions, insulin treatment resulted in decreased VAPB-Nup358 interaction, but increased VAPB-mTOR association (**Fig. 4c, d**). These results indicate a counteracting role for Nup358 and mTORC2 on the stability of ERMCS during growth factor signalling, possibly mediated by their mutually exclusive interaction with VAPB-PTIPI51 complex at the ERMCS.

### Nup358 depletion induces autophagy

The finding that Nup358, as a component of AL, is present at the ERMCS and regulates their dynamics, prompted us to investigate if it is involved in any specific functions mediated by the ERMCS. Autophagy, a key cellular process regulated at the ERMCS, is a conserved cell survival strategy that enables re-utilisation of the cellular components under unfavourable conditions, and maintenance of cellular homeostasis via removal of damaged organelles and aggregated proteins (Gomez-Suaga et al., 2017; Hamasaki et al., 2013; Morishita and Mizushima, 2019). ERMCS act as a platform for the formation of autophagosomes, a double-layered membrane that engulfs cellular components and delivers them to the lysosomes for degradation (Morishita and Mizushima, 2019).

To investigate if Nup358 is involved in autophagy regulation, the levels of autophagy specific markers were assessed under Nup358 depleted conditions in HeLa and HEK293T cells. siRNA-mediated knockdown of Nup358 in HeLa cells led to increased autophagy, indicated by elevated lipidated LC3 (LC3 II) and Beclin1, and decreased p62 levels (**Fig. 5a**). Increased LC3 II and Beclin1 levels were also detected in Nup358 KO HeLa cells (**Fig. 5b**) as well as in HEK293T cells deficient for Nup358 (**Supplementary Fig. 4a**). Additionally, depletion of Nup358 resulted in an increase in the number of LC3 puncta in HEK293T cells stably expressing GFP-LC3, indicative of increased autophagy levels (**Supplementary Fig. 4b**).

**Fig. 5.**
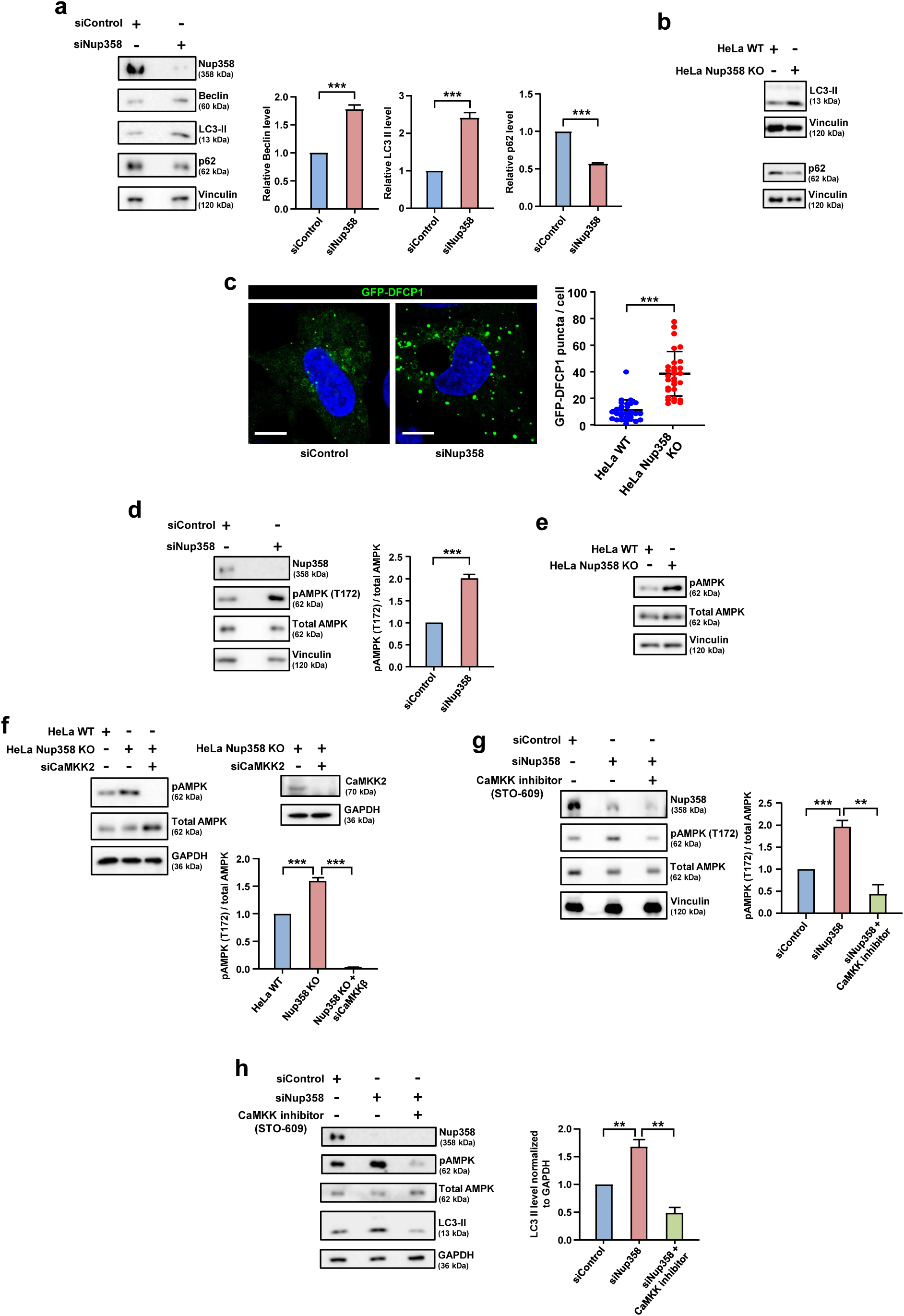
Nup358 deficiency induces autophagy through CaMKK2-mediated activation of AMPK. **a**, Depletion of Nup358 induces autophagy. Hela cells were treated with siControl or siNup358 and monitored for autophagy by analysing the levels of specific proteins by western blotting. Left: Extent of Nup358 depletion and levels of different autophagy markers were examined using specific antibodies. Vinculin was used as loading control. Right: quantitative data showing the changes in levels of shown proteins (n = 3 independent experiments). Data are mean±SEM, unpaired Student’s *t* test. *** *P*≤0.001. **b**, Induction of autophagy was monitored in wild-type (WT) and Nup358 knockout (KO) HeLa cells using indicated antibodies. Vinculin was used as loading control. **c**, Nup358 depletion increases autophagic flux. HeLa cells were initially transfected siControl or siNup358 and later with GFP-DFCP1 and the extent of DFCP1 puncta formation was analysed. Left: representative fluorescent image showing the number of puncta (green) under the described condition. Scale bar, 10 μm. Right: quantitative data depicting the number of GFP-DFCP1 puncta per cell under siControl and siNup358 conditions (*n* = 30 cells from one experiment). Data are mean±SEM, unpaired Student’s *t* test. *** *P*≤0.001. **d**, Nup358 depletion activates AMPK. HeLa cells treated with siControl or siNup358 were analysed for the activation of AMPK based on the status of AMPK phosphorylation at T172. Left: Nup358 depletion and levels of modified and total proteins were monitored by western blotting using indicated antibodies. Right: quantitative data showing the extent of AMPK activation in the presence (siControl) or absence (siNup358) of Nup358 (*n* = 3 independent experiments). Data are mean±SEM, unpaired Student’s *t* test. *** *P*≤0.001. **e**, Activation of AMPK in Nup358 KO HeLa cells as compared to WT cells, assessed by western blotting using specific antibodies. Vinculin was used as loading control. **f**, Co-depletion of CaMKK2 reverses the activation of AMPK caused due to Nup358 depletion. WT and Nup358 KO HeLa cells were treated with specific siRNAs as shown and analysed for the activation status of AMPK. Top left: Extent of AMPK activation under different conditions was assessed by pAMPK (T172) antibody. Top right: Western blotting showing the depletion of CaMKK2. GAPDH was used as loading control. Bottom: quantitative representation of relative pAMPK levels under the described conditions (*n* = 3 independent experiments). Data are mean±SEM, unpaired Student’s *t* test. *** *P*≤0.001. **g**, Chemical inhibition of CaMKK2 using STO-609 reverses the AMPK activation due to Nup358 depletion. HeLa cells were transfected with siRNAs and treated with STO-609 (+) or vehicle control (-) as indicated. Left: cells were analysed for AMPK activation using pAMPK (T172) specific antibody. Vinculin was used as loading control. Right: Quantitative data depicting the relative levels of pAMPK as compared to total AMPK under the described conditions (*n* = 3 independent experiments). Data are mean±SEM, unpaired Student’s *t* test. ** *P*≤0.01, *** *P*≤0.001. **h**, Activation of AMPK and induction of autophagy in Nup358-deficient cells are dependent on CaMKK2. HeLa cells treated with siControl or siNup358 in the presence of vehicle control (-) or CaMKK2 inhibitor STO-609 and monitored for AMPK activation status and induction of autophagy by western blotting using specific antibodies (left). GAPDH was used as loading control. Right: Relative levels of LC3-II as compared to GAPDH under indicated conditions were quantitated (*n* = 3 independent experiments). Data are mean±SEM, unpaired Student’s *t* test. ** *P*≤0.01.

Enhanced autophagy observed under Nup358 depletion could be due to an increase in the autophagic flux or compromised fusion of autophagosome with lysosomes (Klionsky et al., 2021). Recruitment of DFCP1 to the autophagosome initiation site is an indicator of autophagic flux, as DFCP1 is not retained in the mature autophagosomes (Axe et al., 2008). Exogenously expressed GFP-DFCP1 formed significantly higher number of puncta in Nup35-deficient cells (**Fig. 5c**). Moreover, absence of Nup358 also led to a decrease in the level of p62 (Fig. 5a, b), an adapter protein that is degraded with the autophagosomes in the lysosome (Mizushima et al., 2010). These results are consistent with the idea of an increased autophagic flux in the absence of Nup358, thus suggesting that Nup358 negatively regulates the autophagy induction.

### Induction of autophagy upon Nup358 loss is mediated by CaMKK2 and AMPK

Two upstream regulators of autophagy are mTORC1 and AMPK; while mTORC1 inhibits autophagy, AMPK activates it (González et al., 2020; Saikia and Joseph, 2021). Localized pool of mTORC1 and AMPK on lysosomes has been shown to regulate energy-stress mediated autophagy (Lin and Hardie, 2018). However, any role for AMPK at the ERMCS in autophagy induction is unclear. Our studies show that loss of Nup358, indeed, activates mTORC1 (**Fig. 2d**), ruling out the possibility that mTORC1 induces autophagy under Nup358 depleted conditions, thus shifting our focus to AMPK. Interestingly, Nup358 depletion in HeLa cells led to enhanced phosphorylation of AMPK at T172, indicative of its activation (González et al., 2020)(**Fig. 5d, e**). Two of the kinases known to stimulate AMPK activity through phosphorylation at Thr172 residue are liver specific kinase (LKB1) and Ca^2+^-dependent kinase kinase 2 (CaMKK2) (Carling et al., 2008; Saikia and Joseph, 2021). As Nup358 depletion leads to hyperphosphorylation of AMPK in the LKB1-deficient HeLa cells, we reasoned that CaMKK2 might be involved in AMPK activation. Consistent with this, depletion of CaMKK2 reversed the hyperactivation of AMPK in Nup358 KO cells (**Fig. 5f**). Moreover, enhanced activation of AMPK and increased autophagy caused by loss of Nup358 could be rescued by chemical inhibition of CaMKK2 (STO-609) (**Fig. 5g, h**). Further, autophagy induction in Nup358-deficient cells could be rescued by chemical inhibition of ULK1, an important kinase involved in the initiation of autophagosomes (**Supplementary Fig. 4c**). Taken together, these results show that Nup358 negatively regulates autophagy by suppressing the CaMKK2/AMPK/ULK1 pathway.

### Increased levels of cytoplasmic Ca^2+^ trigger AMPK activation in Nup358-deficient cells

As CaMKK2 is a Ca^2+^-induced kinase (Swulius and Waxham, 2008), we reasoned that Nup358 depletion led to an increase in cytoplasmic Ca^2+^, which in turn activated CaMKK2 and AMPK. The Ca^2+^ indicator dye Fluo3-AM, revealed a remarkable increase in the cytoplasmic Ca^2+^ levels when Nup358 was depleted (**Fig. 6a**). BAPTA-AM mediated chelation of cytoplasmic Ca^2+^ was sufficient to rescue AMPK hyperactivation in Nup358-deficient cells (**Fig. 6b**). Collectively, the data shows that Nup358 negatively regulates AMPK activation by limiting the cytoplasmic Ca^2+^ level.

**Fig. 6.**
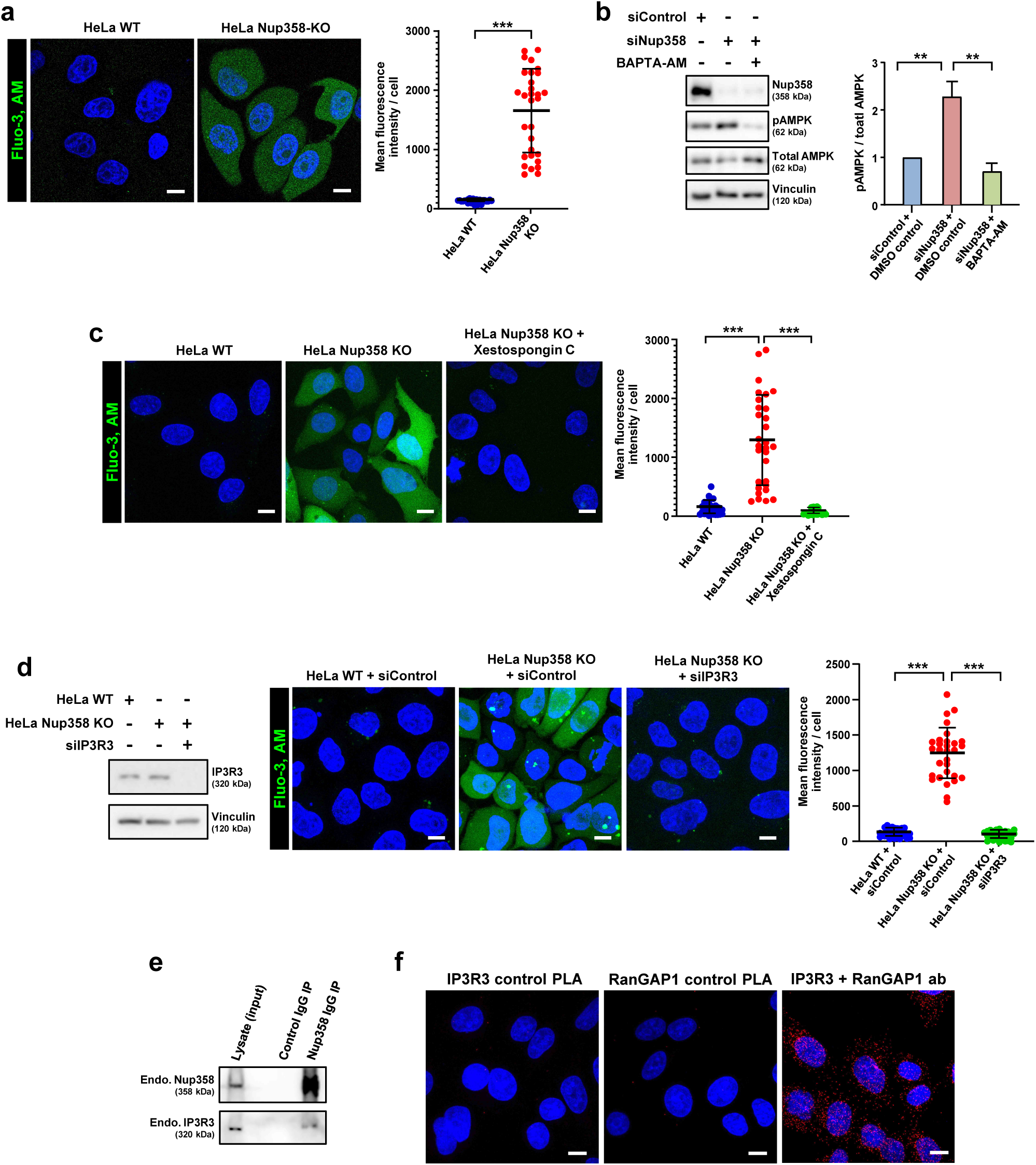
Nup358 interacts with IP3R to restrict release of Ca^2+^ into the cytoplasm and inhibit AMPK activation. **a**, Nup358 depletion leads to Ca^2+^ accumulation in the cytoplasm. WT and Nup358 KO HeLa cells were incubated with Fluo-3, AM (4 μM, green) for 15 min and the fluorescence was monitored by confocal microscopy. Left: Representative microscopic images showing calcium accumulation in the Nup358-deficicent (Nup358 KO) cells as compared to the WT HeLa cells. Scale bar, 10 μm. Right: quantitative data depicting the mean fluorescence intensity of each cell derived from WT and Nup358 KO cells as shown (*n* = 30 cells from 3 independent cells). Data are mean±SD, Mann-Whitney test. *** *P*≤0.001. **b**, Chelation of cytoplasmic Ca^2+^ reverses the activation of AMPK in Nup358-deficient cells. HeLa cells treated with siControl or siNup358 were incubated with vehicle control (-) or BAPTA (+) (20 μM) for 1 h. Left: Western analysis to monitor the depletion of Nup358 and activation status of AMPK using specific antibodies. Vinculin was used as loading control. Right: Quantitative representation of relative pAMPK levels under indicated conditions (*n* = 3 independent experiments). Data are mean±SEM, unpaired Student’s *t* test. ** *P*≤0.01. **c**, Chemical inhibition of IP3R reverses the cytoplasmic accumulation of Ca2^+^ in Nup358 depleted cells. HeLa WT and Nup358 KO cells were treated with vehicle control (-) or Xestospongin C (+) (4 μM for 15 min) under the mentioned conditions and were incubated with Fluo-3, AM (4 μM) for 15 min. Left: representative images showing Fluo-3, AM fluorescence (green). Scale bar, 10 μm. Right: Quantitative depiction of mean fluorescence intensity per cell derived from indicated conditions (*n* = 30 cells from 3 independent experiments). Data are mean±SD, Mann-Whitney test. *** *P*≤0.001. **d**, Co-depletion of IP3R3 reverses cytoplasmic accumulation of Ca^2+^ in Nup358-deficient cells. WT and Nup358 KO HeLa cells were treated with specific siRNAs and cytoplasmic Ca^2+^ accumulation was assessed using Fluo-3, AM. Left: Western blots showing the extent of depletion of IP3R3. Vinculin was used as loading control. Middle: representative microscopic images displaying Fluo-3, AM fluorescence (green). Scale bar, 10 μm. Right: Quantitative data depicting mean fluorescence intensity per cell derived from cells under the described conditions (*n* = 30 cells from 3 independent experiments). Data are mean±SD, Mann-Whitney test. *** *P*≤0.001. **e**, Nup358 interacts with IP3R3. Endogenous Nup358 was immunoprecipitated from HeLa cells and presence of IP3R3 in the IP samples were analysed by western blotting. **f**, In situ PLA confirmed the interaction between endogenous Nup358 and IP3R3. In this assay, RanGAP1, another protein that tightly associates with Nup358 (Joseph et al., 2004), was used instead of Nup358. IP3R3 or RanGAP1 antibody alone was used as control for PLA. Scale bar, 10 μm.

### Nup358 interacts with IP3R and restricts Ca^2+^ release from the ER

IP3 receptor (IP3R) is a channel that release Ca^2+^ into the cytoplasm from its major store house – the ER (Prole and Taylor, 2019). Moreover, IP3R is enriched at the ERMCS (Csordás et al., 2018; Grimm, 2012), where Nup358 is also present. We therefore investigated a plausible role for IP3R in Nup358 mediated regulation of cytoplasmic Ca^2+^ levels. Consistent with this idea, treatment with Xestospongin C, a potent inhibitor of IP3R, rescued the accumulation of cytoplasmic Ca^2+^ in Nup358 KO HeLa cells (**Fig. 6c**). These findings were confirmed by siRNA-mediated depletion of IP3R3 (the major isoform of IP3R in HeLa cells) in Nup358 KO cells (**Fig. 6d**). The data thus suggests that the elevation of cytoplasmic Ca^2+^ levels in Nup358-deficient cells depends on IP3R mediated Ca^2+^ release. Moreover, co-immunoprecipitation assays revealed an interaction between endogenous Nup358 with IP3R3 (**Fig. 6e**), again confirmed by PLA (**Fig. 6f**). Collectively, the data implies a key role for Nup358 in restricting autophagy through the IP3R/Ca^2+^/CaMKK2/AMPK axis.

Recently, a role for AMPK in the activation of mTORC2 has been elucidated (Kazyken et al., 2019), opening up the possibility that increased cytoplasmic Ca^2+^ triggers AMPK activation, which subsequently may activate mTORC2 in Nup358-deficient cells. Co-depletion of AMPK and Nup358 was performed to verify the same. However, AMPK depletion did not rescue the activation of mTORC2/Akt in Nup358 knockdown cells (**Supplementary Fig. 5a**). Simultaneously, we also investigated if mTORC2/Akt could activate AMPK under our experimental conditions. Disrupting mTORC2 activity by depletion of Rictor did not reverse the AMPK activation upon Nup358 knockdown (**Supplementary Fig. 5b**). Taken together, these data indicate that restriction of mTORC2/Akt and AMPK activity by Nup358 may occur through independent mechanisms.

### A conserved role for Nup358 in restricting mTORC2 activity, cytoplasmic Ca^2+^ levels and autophagy in Drosophila

Our studies uncovered a function for Nup358 in regulating mTORC2/Akt dependent ERMCS stability and Ca^2+^-induced autophagy in mammalian cells. The human and Drosophila Nup358 (dNup358) proteins have a significant degree of conservation in their domains (Forler MCB 2004). We, therefore, investigated if the novel functions of Nup358, elucidated above, are conserved in Drosophila. Interestingly, inducible RNAi-dependent loss of Nup358 in Drosophila (**Fig. 7a**) augmented the phosphorylation of Akt at S505 (equivalent of S473 in humans) (**Fig. 7b**) and AMPK at T184 (equivalent of T172 in humans) (**Fig. 7c**). These results indicate that Nup358’s role in restricting mTORC2/Akt and AMPK activity is functionally conserved between humans and flies.

**Fig. 7.**
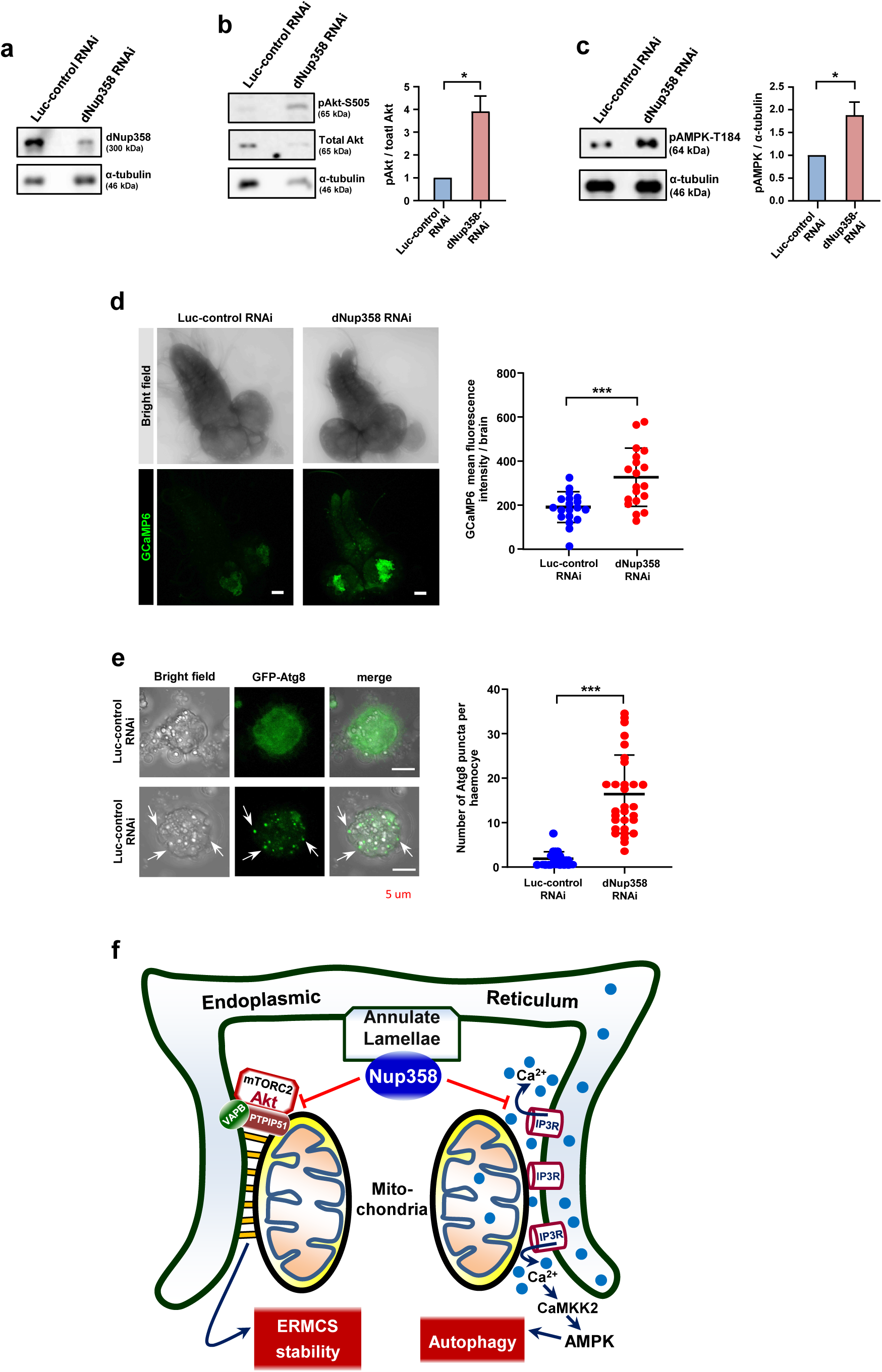
Conserved regulatory role of Nup358 in restricting mTORC2/Akt activation, AMPK stimulation, cytoplasmic Ca^2+^ level and autophagy in Drosophila. a, b, c, Knockdown of Nup358 from Drosophila as assessed by western blotting. Adult flies were treated with 500 μM mifepristone (RU486) for 72 h to induce shRNA for control (Luciferase, Luc) or Nup358. Brains lysates were assessed for dNup358 knockdown by western blotting. Tubulin was used as loading control (**a**). The lysates were analysed for pAkt (S505) (**b**, left) and pAMPK (**c**, left) levels by western blotting. (**b, c**, right) quantitative data showing the levels of pAkt and pAMPK under indicated conditions (*n* = 3 independent experiments). Data are mean±SEM, unpaired Student’s *t* test. * *P*≤0.05. **d**, Nup358 knockdown in Drosophila leads to accumulation of cytoplasmic Ca^2+^ in the larval brain. GCaMP6-expressing Luc-control or dNup358 RNAi lines (L3 larval stage) were monitored for GCaMP6 fluorescence. Left: Representative microscopic images displaying GCaMP6 fluorescence (green). Scale bar, 50 μm. Right: Quantitative data depicting the mean fluorescence intensity per brain derived from Luc-control and dNup358 RNAi lines (*n* = 19 brains from 3 independent experiments). Data are mean±SD, Mann-Whitney test. *** *P*≤0.001. **e**, Nup358 deficiency leads to increased autophagy in Drosophila. Drosophila haemocytes derived from GFP-Atg8 expressing Luc-control or Nup358 RNAi larvae were analysed for Atg8 puncta formation. Left: Show the representative microscopic images obtained under the mentioned conditions. Right: Quantitative data depicting the number of GFP-Atg8 puncta per haemocyte derived from Luc-control and dNup358 RNAi lines (*n* = 30 cells from 3 independent experiments). Data are mean±SD, Mann-Whitney test. *** *P*≤0.001. **f**, Model depicting the function of AL-resident Nup358 in regulating ERMCS functions. AL-associated Nup358 resides at the ERMCS. Our studies revealed a role for the ERMCS tethering complex VAPB-PTPIP51 in binding and activating mTORC2/Akt at the ERMCS upon growth factor signalling, consequently strengthening the contacts between these two organelles. Our data also suggest that both Nup358 and mTORC2 have a reciprocal binding relationship with VAPB-PTPIP51 complex, which ultimately determines the extent of ERMCS stability. Previous studies have reported that Ca^2+^-stimulated autophagy is mediated through CaMKK2 and AMPK (Saikia and Joseph, 2021; Sun et al., 2016). Our results reveal an important role for Nup358 in binding to the ERMCS-enriched Ca^2+^ channel IP3R and limiting its ability to release Ca^2+^ into the cytoplasm, thereby inhibiting Ca^2+^-triggered autophagy.

Further, to test if dNup358 restricted the cytoplasmic Ca^2+^ level, Nup358 was knocked down in the Drosophila brain using a pan-neuronal expressing Elav Gal4 line that also expressed the cytoplasmic Ca^2+^ reporter GCaMP6. Compared to the Luc RNAi control, loss of Nup358 led to an enhanced accumulation of Ca^2+^ in the brain (**Fig. 7d**), indicating the functional conservation of Nup358 in limiting the cytoplasmic Ca^2+^ levels. Furthermore, Nup358 knockdown in Drosophila haemocytes expressing GFP-LC3 (Atg8a) led to enhanced autophagy as revealed by increased LC3 puncta (**Fig. 7e**), supporting a conserved role for Nup358 in negatively regulating autophagy in the flies.

Based on the results, we propose the following working model (**Fig. 7f**). Nup358, as a part of AL, resides at the ERMCS wherein it regulates their dynamics and associated functions for the maintenance of cellular homeostasis. We propose that upon growth factor stimulation, an initial recruitment of mTORC2/Akt to the ERMCS occurs via its interaction with the VAPB-PTIPI51 complex. This interaction results in growth factor-dependent activation of mTORC2 and a subsequent stabilization of ERMCS mediated by Akt-dependent phosphorylation of ERMCS proteins such as PACS2 (Betz et al., 2013). The recruitment of mTORC2/Akt, its activity and the stability of ERMCS are restricted by AL-resident Nup358, thereby controlling the overall ERMCS dynamics. Nup358 also limits cytoplasmic Ca^2+^ release by IP3R and inhibits Ca^2+^-induced autophagy, possibly via a mechanism independent of its effect on mTORC2 and/or ERMCS dynamics.

## DISCUSSION

Our studies provide new insights into the cellular function of the underexplored organelle AL. The results show that Nup358-labelled AL are often present at the contact sites between ER and mitochondria, which are known to play a pivotal role in cellular homeostasis by regulating critical processes such as inter-organelle transfer of Ca^2+^ and lipids, mitochondrial energetics, cellular metabolism, apoptosis and autophagy. Depletion of Nup358 led to increased ERMCS stability through hyperactivation of mTORC2/Akt, and increased autophagy mediated through stimulation of Ca^2+^/CaMKK2/AMPK/ULK1 pathway. Thus, the findings of our study highlight the importance of AL-resident Nup358 for achieving cellular homeostasis by modulating the ERMCS dynamics and functions.

While investigating the role of Nup358-positive AL in cellular functions, we identified a mechanism by which ERMCS components regulate mTORC2/Akt activation in response to growth factor signalling. Earlier studies have shown that growth factors can stimulate the recruitment of mTORC2/Akt to ERMCS, leading to Akt-mediated phosphorylation of proteins at the contacts, thereby regulating the dynamics and functions of ERMCS (Betz et al., 2013). The mTORC2/Akt complex has been shown to be present at multiple intra-cellular locations, but the site and mechanism of its activation upon growth factor signalling remain undetermined (Ebner et al., 2017; Fu and Hall, 2020). Our studies indicate that the ERMCS tethering complex VAPB-PTPIP51 is important for the activation of mTORC2/Akt pathway, possibly via its recruitment to the ERMCS. This is consistent with an earlier report that illustrated the significance of the integrity of the ERMCS platform in insulin signalling (Tubbs and Rieusset, 2017; Tubbs et al., 2014). Our studies reveal a molecular interplay between Nup358 and mTORC2/Akt at the ERMCS for the regulation of ERMCS dynamics. A mutually exclusive binding of Nup358 and mTORC2 complex to the ERMCS tethering complex VAPB-PTPIP51 appears to regulate the growth-factor mediated ERMCS dynamics and associated functions. In support of this, earlier studies have shown that Nup358 haploinsufficiency in mice leads to metabolic disturbances and defective glucose homeostasis (Aslanukov et al., 2006). Our finding that Nup358 functions in the fine-tuning of growth factor-dependent activation of mTORC2 pathway will help explore the detailed mechanism of mTORC2 activation in general and the specific contribution of the ERMCS.

Additionally, our studies highlight the importance of Nup358 in Ca^2+^ homeostasis, specifically by restricting the Ca^2+^ release by the ER-resident channel IP3R. Although we identified an interaction between Nup358 and IP3R, the molecular mechanism by which Nup358 inhibits IP3R function warrants further research. In Nup358-deficient cells, as a consequence of dysfunctional Ca^2+^ homeostasis, AMPK is activated which further triggers autophagy. Although cytosolic Ca^2+^ is known to stimulate autophagy through CaMKK2-mediated phosphorylation of AMPK (Carling et al., 2008), the physiological relevance is unclear. Nup358 mediated regulation of Ca^2+^ release by IP3R might impact other Ca^2+^-mediated functions (Bagur and Hajnóczky, 2017). Nevertheless, our results indicate that the AL-resident Nup358 is a crucial regulator of cellular homeostasis by modulating mTORC2 signalling, ERMCS dynamics, cytoplasmic Ca^2+^ levels and autophagy.

In light of the new findings, it is unclear as to why AL components are involved in regulating cellular signalling and cytoplasmic processes. As the major component of AL identified so far are nucleoporins, which are present on the NPCs as well, it is possible that there exists a cross-talk between nucleo-cytoplasmic transport (NCT) and cytoplasmic signalling events such as nutrient sensing and autophagy. Based on the ultra-structural studies, an interconnection between the nuclear envelope and AL has been proposed (Kessel, 1992). Moreover, budding of nucleoporin-containing structures and fusion of them with pre-existing AL in the cytoplasm have been observed (Sahoo et al., 2017). The possible interconnection between NCT and AL is particularly interesting in the light of the implications of dysregulated NCT on the pathogenesis of neurodegenerative diseases (Ding and Sepehrimanesh, 2021). The newly identified role for AL-resident nucleoporins in regulating events other than NCT might be relevant in the pathophysiology of neurodegenerative diseases.

Nup358 mutations are linked with acute necrotizing encephalopathy 1 (ANE-1), a disease condition triggered by viral infection, and characterized by lesions in specific regions within the brain of affected individuals (Neilson, 2010; Neilson et al., 2009). The pathophysiology includes cytokine storm, exemplified by increased levels of pro-inflammatory cytokines such as INF-α and IL-6 in the serum and cerebrospinal fluid of the patients (Levine et al., 2020). Impairment in the miRNA-mediated translation regulation by Nup358 might contribute to the development of ANE-1 (Deshmukh et al., 2021; Shen et al., 2021). However, the functions for AL-associated Nup358 revealed by our studies may provide an additional platform for a better understanding of the pathogenesis of ANE-1.

## Methods

### Mammalian cell culture and treatments

HeLa S3, HEK293T, U2OS and HuH7 cells were cultured in DMEM (Invitrogen) with 10% FBS and 10 μg/ml ciprofloxacin (Fresenius Kabi, India). Plasmid constructs were transfected using polyethyleneimine, linear (PEI, MW-25,000; Polysciences Inc.) or Lipofectamine 2000 (Invitrogen).

### Chemicals

The following reagents were used in the study. Insulin (I6634, Sigma), EGF (E9644, Sigma) STO-609 acetic acid (S1318, Sigma), Mifepriston-RU486 (M8046, Sigma) were purchased from Sigma. BAPTA, AM (2787, Tocris Bioscience), Xestospongin C (ab120914, Abcam), Fluo-3, AM (F1241, Invitrogen), MRT68921 HCI (S7949, Selleckchem), Pluronic F-127 (Sigma, P2443), MitoTracker Deep Red FM (Invitrogen, M22426).

### Antibodies

The following antibodies were used in this study. Anti-Nup358 (Joseph et al., 2004), (western blot (WB),1:3,000; immunofluorescence (IF); proximity ligation assay (PLA), 1:600), -dNup358 (WB, 1:3,000) and -PTPIP51 (PLA, 1:250) were produced in-house. Anti-mTOR (2983; WB, 1:4,000; PLA, 1:400), -Rictor (2140; PLA,1:100), Mitofusin 2 (9482S; WB, 1:4,000) -LC3A/B (12741; WB, 1:5,000), -Beclin1 (3495; WB, 1:500), -p62 (5114; WB, 1:500), pAMPK-T172 (2535; WB, 1:3,000), AMPK (2793; WB, 1:3000), pAkt-S473 (9271; WB, 1:3,000), pAkt-T308 (9275; WB, 1:3,000), Akt (9272; WB, 1:3,000), pS6K (9205; WB, 1:3,000), S6K (9202; WB, 1:3,000), pS6 (2211; WB 1:4,000) and S6 (2317; WB, 1:4,000) were purchased from Cell Signalling Technology. Anti-IP3R3 (610312; WB, 1:3,000), -Beclin (612112; WB, 1:3,000) -p62/lck (610832; WB, 1:2,000), -Nup88 (6111896; WB, 1:3,000), -Nup62 (610497; WB, 1:4,000) and -Tom20 (612278; IF, 1:200) were purchased from BD biosciences. Anti-PDIA3 (AMAB90988; WB, 1:3000; IF, 1:500), RanGAP1 mouse (33-0800; IF, 1:300; PLA, 1:500) was from Invitrogen. Anti-VAPB (66191-1-1g; WB, 1:4,000), -PTPIP51 (20641-1-AP; WB, 1:3,000), -CaMKK2 (11549-I-AP; WB, 1:200), -LC3 (14600-1-AP; WB, 1:3,000) α-tubulin (66031-1-Ig, WB, 1:5,000) were purchased from Proteintech. Anti-Rictor (ab104838; WB, 1:2,000), AMPK (ab32047; WB, 1:3,000), Nup214 (ab70497; IF, 1:100) were purchased from Abcam. Anti-myc (sc-401; WB, 1000), -Tom20 (sc17764; WB, 1:3,000) and GAPDH (sc-166574; WB, 1:5,000) were from Santa Cruz Biotechnology Inc. Other antibodies used in the studies were Anti-FLAG (F3165; WB, 1:5,000) and -vinculin (V9131; WB, 1:10,000) from Sigma, anti-VAPB (VMA00454; PLA, 1:600) from BioRad, Anti-IP3R3 (07-1213; PLA, 1:500) from Millipore, anti-HA (MMS-101R; WB, 1:5,000) from Covance Research Products (BioLegend), and anti-TNRC6A/GW182 (A302-329A; WB, 1:500) from Bethyl Laboratories.

### DNA Constructs

pCI-neo-HA-PTPIP51 and pCI-neo-myc-VAPB were gifts from Christopher Miller (King’s College London, UK). pCRISPR-CG01 (HCP216100-CG01-1-10) was purchased from GeneCopoeia and pcDNA3-Flag-mTOR wt (ID-26603) was procured from Addgene. Mito-BFP, RFP-KDEL (ER) and pEGFP-LC3 were provided by Jennifer Lippincott-Schwartz (Janelia Research Campus, USA), pEGFP-DFCP1 was from Nicholas Ktistakis (Babraham Institute, UK) and pcDNA3-hPACS2-3HA was from Gary Thomas (University of Pittsburgh, USA).

### siRNAs

The following sequences were used for siRNAs. siNup358 (5′ GGTGAAGATGGATGGAATA 3′) (Sahoo et al., 2017), siControl 5′ TTCTCCGAACGTGTCACGT 3′) (Sahoo et al., 2017), siVAPB (5’ GCTCTTGGCTCTGGTGGTT 3’), siPTPIP51 (5’ CCTTAGACCTTGCTGAGAT 3’), siMfn2 (5’ GTGATGTGGCCCAACTCTA 3’), siRictor (5’ GCAGCCTTGAACTGTTTAA 3’) siCAMKK2B (5′ GCAUCGAGUACUUACACUAUU 3’) were purchased from Dharmacon. siITPR3/IP3R3 (4392420, ID s266), siPACS2 (4392420, ID s23371) and siRNAs against GW182 isoforms, siTNRC6A/GW182 (4390824, ID s26154) and siTNRC6B (4392420, IDs 23060), were obtained from ThermoFisher Scientific (Ambion). siAMPK α1/2 (sc-45312) was purchased from Santa Cruz Biotechnology Inc. All siRNA transfections (40 nM, 72 h) were performed using Lipofectamine RNAiMAX (Invitrogen) as per manufacturer’s instruction.

### Co-immunoprecipitation (co-IP) and western blot (WB) analysis

In general, cell lysates for immunoblot analysis were prepared in 1% NP40 buffer contain 20 mM Tris-HCl (pH 8), 137 mM NaCl, 10% glycerol, 2 mM EDTA, supplemented with 7.5 mM NaF, 0.75 mM sodium orthovanadate, 1 mM PMSF, protease inhibitor cocktail of leupeptin (50 μg/ml), aprotinin (5 μg/ml), and pepstatin (2 μg/ml). After thorough retro pipetting, cell debris was pelleted down by spinning at 12,900*g* for 10 min. A clear supernatant was collected and protein concentration was estimated using Bradford (Bio-Rad) assay. Samples were subjected to SDS-PAGE and transferred to PVDF membrane (Millipore) for western analysis.

For extracting phosphoproteins, cell lysis was performed in a buffer containing 50 mM Tris-HCl (pH 7.5), 1 mM EDTA, 1% Triton X-100, 10% glycerol, 1 mM DTT, 50 mM sodium fluoride, 5 mM sodium pyrophosphate, 1 mM PMSF supplemented with protease inhibitor cocktail (Roche).

For immunoprecipitation of endogenous PTPIP51 or Nup358, the cell extracts were made in 1% NP40 cell lysis buffer as previously described. The whole cell extracts were incubated with Protein A sepharose beads (Invitrogen) coated with anti-PTPIP51 or anti-Nup358 antibody for 2 h at 4 °C. The precipitates were washed once with 1% NP40 cell lysis buffer, followed by two washes with tris-buffered saline (TBS). The IP samples were subjected to SDS-PAGE and analysed by WB.

For IP of mTOR, cells were lysed in 0.5% CHAPS buffer containing 120 mM NaCl, 40 mM HEPES (pH 7.5), 0.2 mM EDTA, 10% glycerol, and FLAG-mTOR was immunoprecipitated with anti-FLAG beads (EZview™ Red ANTI-FLAG^®^ M2 Affinity Gel, Sigma, F3165). The immunoprecipitates were washed three times with 0.5% CHAPS buffer before proceeding with WB.

For the interaction studies involving Nup358 and IP3R3, cells were lysed in 1% CHAPS buffer containing 150 mM NaCl, 50 mM Tris-HCl (pH 7.4), and incubated with anti-Nup358 antibody coated protein A/G sepharose beads for 2 h at 4 °C. Immunoprecipitates were further washed with CHAPS buffer followed by TBS and analysed by WB.

### Subcellular fractionation

ERMCS (MAM) fractionation was performed from HeLaS3 cells based on the standardized MAM isolation protocol (Wieckowski et al., 2009; Williamson et al., 2015). Cells from six T-175 confluent flasks were trypsinized and pelleted by centrifugation at 600*g* for 5 min. All the further steps were performed at 4°C using fractionation buffers supplemented with protease inhibitor cocktail. The cell pellet was suspended in IB-1 buffer [225 mM mannitol, 75 mM sucrose, 0.1 mM EGTA and 30 mM Tris-HCl (pH 7.4)] and homogenized by passing through syringes of different nozzle sizes (21Ga, 23Ga and 24Ga), 10 times each. Post nuclear fraction (PNF) was collected from the homogenate by centrifugation at 600*g* for 5 min. Later, for pelleting crude mitochondria, the PNF was centrifuged at 7,000*g* for 10 min and the supernatant containing soluble proteins and ER/microsomes was collected as cytoplasmic fraction. To avoid the microsomal contamination, the crude mitochondrial pellet was resuspended in IB-2 buffer [225 mM mannitol, 75 mM sucrose, and 30 mM Tris-HCl (pH 7.4)] and centrifuged at 7,000*g* for 10 min. This step was repeated once again, centrifuging at 10,000*g*, this turn. The final pellets were washed and reconstituted in MRB buffer [250 mM mannitol, 5 mM HEPES (pH 7.4), and 0.5 mM EGTA)] and gently layered on top of the percoll medium [225 mM mannitol, 25 mM HEPES (pH 7.4), 1 mM EGTA, and 30% percoll]. After 30 min of centrifugation at 95,000*g*, the ERMCS (MAM) fraction, a whitish diffused band at the midst of centrifuge tube, was carefully collected with a Pasteur pipette. The remaining lower layer, containing mitochondrial suspension, was diluted in MRB solution, and mitochondria were extracted by centrifugation at 6,300*g* for 20 min. Similarly, the ERMCS fraction was resuspended in MRB buffer and centrifuged at 100,000*g* for 60 min to obtain the ERMCS pellet. All the final fractions were dissolved in RIPA buffer [50 mM Tris-HCl (pH 7.4), 150 mM NaCl, 1% NP-40, 0.1% SDS, 0.5% sodium deoxycholate] and analysed by western blotting.

### Immunofluorescence (IF) and Proximity ligation assay (PLA)

Cells seeded on coverslips were fixed using methanol (−20°C) for 5 min. After washing with phosphate-buffered saline (PBS), cells were incubated in primary antibodies diluted in 2% normal horse serum (NHS, Vector Laboratories) for 60 min. After a PBS wash, Alexa Fluor labelled secondary antibodies (Molecular probes, ThermoFisher Scientific) diluted in 2% NHS were added and incubated for 30 min. For staining of the nucleus, Hoechst 33342 dye (Sigma) was added to the secondary antibody solution. After three washes with PBS, the coverslips were mounted with Vectashield medium (Vector Laboratories). Images were acquired using Olympus FV3000 confocal laser scanning microscope with the 60x, 1.42 NA or 100x, 1.45 objective. The images were processed with cellSens (ver.2.3) constrained iterative deconvolution for better resolution of ER and mitochondrial network.

Proximity Ligation assay (PLA) was used to identify *in situ* protein interactions. Briefly, cells grown on coverslips were fixed with methanol (−20°C) for 5 min and probed with the respective primary antibodies. PLA was performed using Duolink reagents as described in Duolink^®^ PLA Fluorescence Protocol (Sigma). The PLA signals developed by Duolink^®^ In Situ Detection Reagents Red (DUO92101) were detected and acquired by Olympus FV 3000 microscope and quantitated using NIH ImageJ.

### Stimulated emission depletion microscopy (STED)

Huh-7 cells grown on high precision microscope cover glasses (Marienfeld, Germany) were fixed with 4% paraformaldehyde (20 min), followed by permeabilization with 0.1% Triton X-100 (10 min). The cells were then incubated with primary antibodies diluted in 2% NHS. After washing the coverslips thrice with PBS, secondary antibodies (Alexa Flour 568 and Alexa Flour 657) prepared in 2% NHS were added, washed thrice with PBS and the coverslips were mounted in Prolong Gold (Invitrogen). Images were acquired using Leica TCS SP8 STED 3X with the 100x, 1.4 NA objective.

### Quantitation of ER and mitochondria colocalization

RFP-ER and Mito-BFP plasmids were co-expressed to visualize the contacts between ER and mitochondria in live cells. Images were analysed for colocalization between ER and mitochondria using ImageJ JACoP software. Manders’ coefficient was calculated from the fraction of mitochondria overlapping with the ER. To visualize the overlapping pixels, represented as ERMCS, ImageJ Colocalization Finder was used.

### Generation of GFP-LC3 stable and Nup358 knockout (KO) cell lines

HEK293T cells were co-transfected with linearized pEGFP-LC3 and pTSiN-puro-Cre (Jan van Deursen Lab, Mayo Clinic, USA) constructs for 72 h. Cells were then selected for puromycin resistance. From the puromycin resistant cells, the GFP-LC3 positive cells were obtained by FACS.

HeLa Nup358 KO cells were generated using CRISPR-Cas9 technology. HeLa cells were co-transfected with Nup358 CRISPR-Cas9 (pCRISPR-CG01, GeneCopoeia) and pTSiN-puro-Cre constructs. Transfected cells were selected for puromycin resistance. Single cell clones were obtained, which were verified for Nup358 KO by IF and immunoblotting. The sgRNA target region within the *Nup358* gene was amplified by PCR, which was sequenced to identify the genomic manipulation occurred (**Supplementary Fig. 3**).

### Ca^2+^ measurement using Fluo-3, AM

Cells seeded on coverslips were incubated with Ca^2+^ free medium for 4 h before treating with 4 μM Fluo-3, AM along with 0.02% Pluronic F-127 for 15 min. Cells were washed with PBS thrice and fixed with 4% paraformaldehyde. Images were acquired immediately. In case of experiments using IP3R inhibitor, cells were initially treated with Xestospongin C (4 μM) for 45 min, later Fluo-3, AM was added to the same medium and incubated for additional 15 min.

### Treatments

In general, prior to western analysis, cells were replenished with fresh medium containing 10% FBS for 24 h. In case when growth factor (insulin and EGF) signalling was assessed under siRNA treated conditions, cells were initially transfected with siRNA for 69 h and the old medium was replaced with serum-free DMEM for 3h. For monitoring the levels of proteins such as LC3, pAMPK and AMPK in HeLa WT and Nup358 KO cells, 40,000 cells per well were seeded in a 24-well plate for 24 h, and the old medium was changed with fresh 10% FBS containing DMEM every 6 h thrice, followed by 90 min and 15 min before cell lysis. For examining p62 levels, Nup358 KO cells were seeded for 24 h as mentioned above and the new medium was added 4 h before lysis. Lysates were further processed for western analysis.

### Antibody production

Antibody against human PTPIP51 was raised in rabbits. Bacterially expressed and purified His-tagged PTPIP51 fragment corresponding to amino acids 244 to 470 was used as the antigen. Specific antibodies were affinity purified using the antigen and used in this study. Antibodies for Drosophila Nup358 were generated in mice against a region corresponding to amino acids 2319 to 2555, expressed and purified from bacteria.

### Drosophila strains and rearing

Flies were maintained on standard Cornmeal Agar at 25 °C and 65% relative humidity in a 12 h light: dark cycle. Gal4/UAS system was used to induce the expression of our gene of interest. Fly strains used in our experiments were Cg-Gal4 (BDSC: 7011), Elav-Gal4 (BDSC: 458), Elav-Gene Switch (BDSC: 43642), UAS-Nup358 RNAi (BDSC: 34967), UAS-Luc Val10 RNAi (BDSC: 35788), UAS-GFP Atg8a (BDSC: 51656), and UAS-GCamp6 (BDSC: 42746).

### Drosophila haemocyte imaging

Flies of Cg-Gal4 and UAS-GFP Atg8a were crossed. Upon eclosion of the F1 progeny, virgin females selected based on GFP expression, were crossed with UAS-Nup358 RNAi or UAS-Luc Val10 RNAi males. L2 and L3 larvae positive for fluorescence were selected and dissections were performed. The haemolymph was collected in 1x PBS, allowed to settle for 10 min, and then used for haemocyte imaging.

### Drosophila GCaMP6 imaging

Flies of Elav-Gal4 and UAS-GCamp6 were crossed. F1 progeny were collected, and virgin females were then crossed with UAS-Nup358 RNAi or UAS-Luc Val10 RNAi males. L3 larvae were picked up based on fluorescence and brains were dissected. Imaging was done in PBS without fixing.

### Drosophila gene switch expression

F1 progeny from the cross of Elav-Gene Switch with either UAS-Nup358 RNAi or UAS-Luc Val10 RNAi were collected upon eclosion. The progeny were starved overnight in vials containing moist paper and fed 500 μM mifepristone (RU486) dissolved in 80% ethanol for 72 h post starvation. Later, the fly heads (*n* = 5) were chopped and lysed in S2 buffer [100 mM NaCl, 50 mM Tris-HCl (pH 7.5), 0.1 % NP40 with protease inhibitors] and processed for western blotting.

### Statistics and reproducibility

All statistical analyses were done in GraphPad Prism version 8, calculated from independently repeated experiments as indicated in the respective Figure legends. For western blot analysis, values were considered from at least three independent experiments (*n* = 3) and a two-sided unpaired Student’s *t* test was performed. The values were graphically expressed as mean ± SEM and *P* values were as indicated in legends. All immunostaining data was analysed by Mann-Whitney U test. The mean ± SD was schematically represented and the level of significance is as stated in the figure legends. In all plots, *P* values are as indicated; *** *P*≤0.001, ** *P*≤0.01 and * *P*≤0.05.

## Acknowledgements

We thank C. Miller, J. Lippincott-Schwartz, N. Ktistakis and G. Thomas for sharing reagents; NCCS Bio-Imaging facility staff for help with microscopy; Richa Ricky (IISER, Pune) and Lizanne Oliveira for critical comments on the manuscript; and members of Joseph and Seshadri Lab for insightful discussions. Stocks obtained from the Bloomington Drosophila Stock Centre (NIH P40OD018537) were used in this study. We thank Girish Ratnaparkhi, IISER, Pune, for sharing UAS-Nup358 RNAi line and Swagatika Panigrahi for the help in generating dNup358 antibody. This work was supported by funding from the Department of Biotechnology (DBT), Ministry of Science and Technology, India, through a grant to J.J. (BT/PR27451/BRB/10/1655/2018) and intramural funding from NCCS. Financial support from the Council of Scientific and Industrial Research, Ministry of Science and Technology, Government of India, to M.K.R. and from DBT to R.S. and P.V. through research fellowships is gratefully acknowledged.

## Author contributions

M.K.R. and J.J. conceived the study. M.K.R., R.S., P.V. A.M. and J.J. designed experiments. M.K.R., R.S., P.V. and P.D. performed experiments. A.M. provided advice on experimental design and data analysis involving Drosophila. M.K.R., R.S. and J.J. wrote the manuscript. All the authors reviewed and edited the manuscript.

