## Supplementary Figures for "Nup358 regulates remodelling of ER-mitochondrial contact sites and autophagy"

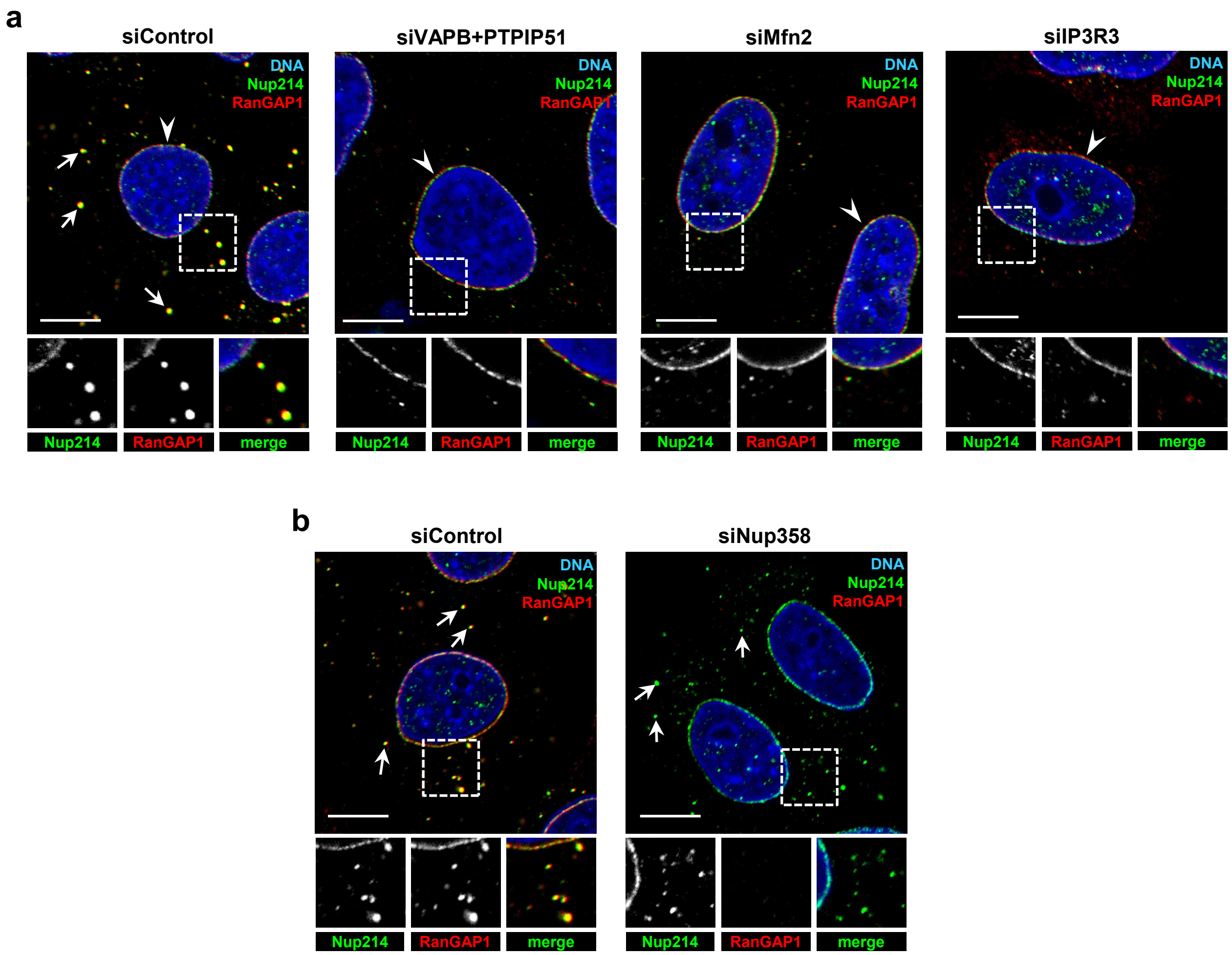


**Supplementary Fig. 1 | ERMCS integrity is important for AL stability. a**, Depletion of ERMCS proteins affects AL assembly. HeLa cells were treated with indicated siRNAs and the AL integrity was monitored by presence of cytoplasmic Nup214 (green) and RanGAP1 (a binding partner of Nup358, red). Number of cytoplasmic puncta, which represent AL (shown in arrows), are significantly reduced when ERMCS proteins were depleted as compared to control cells. Under the same conditions, the NE staining of nucleoporins (arrowheads) was largely unaffected. Scale bar, 10 μm. **b**, Depletion of Nup358 does not affect AL integrity. HeLa cells were treated with siControl or siNup358 and were immunostained for Nup214 (as an AL marker, green) and RanGAP1 (red). Note that Nup214 positive AL (arrows) puncta largely remained unaffected. Scale bar, 10 μm.

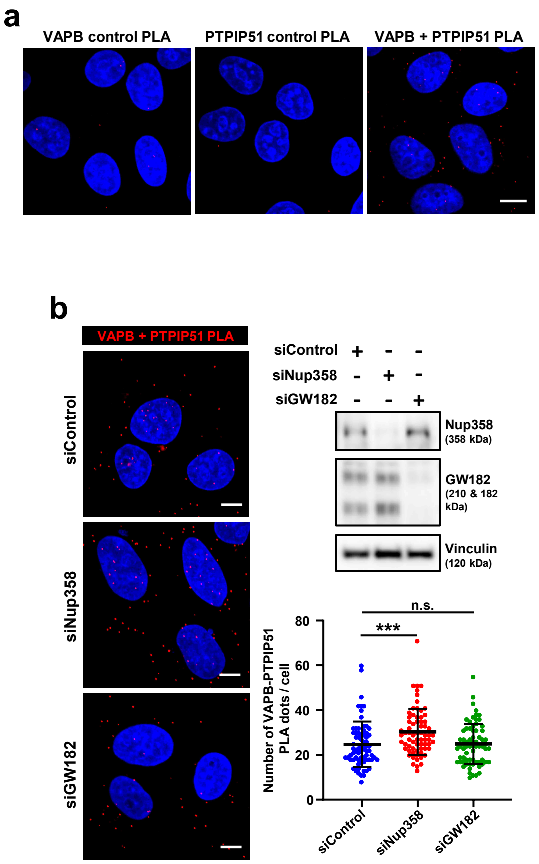


**Supplementary Fig. 2 | Interfering with miRNA pathway does not affect ERMCS integrity. a**, Specificity of in situ PLA with VAPB and PTPIP51 antibodies. HeLa cells were either incubated with antibodies against VAPB (control PLA) or PTPIP51 (control PLA) or both (VAPB+PTPIP51 PLA) and proceeded with PLA. Specific PLA puncta (red) were apparent when incubated with both antibodies (VAPB+PTPIP51) as compared to single antibody control. Scale bar, 10 µm. **b**, Depletion of GW182 does not affect contacts between ER and mitochondria. HeLa cells were treated with indicated siRNAs and the intactness of ERMCS was monitored *in situ* by PLA using VAPB and PTPIP51 antibodies. Left: Depletion of indicated proteins were confirmed by western blotting. Vinculin was used as loading control. Middle: Representative microscopic images showing PLA puncta (red) under indicated conditions. Scale bar, 10 µm. Right: quantitative data showing number of PLA dots per cell, derived from indicated conditions ($n$ = 70 cells for siControl; 69 cells for siNup358 and 71 cells for siGW182 from 3 independent experiments). Data are mean±SD, Mann-Whitney test. n.s. not significant, *** $P \leq 0.001$.

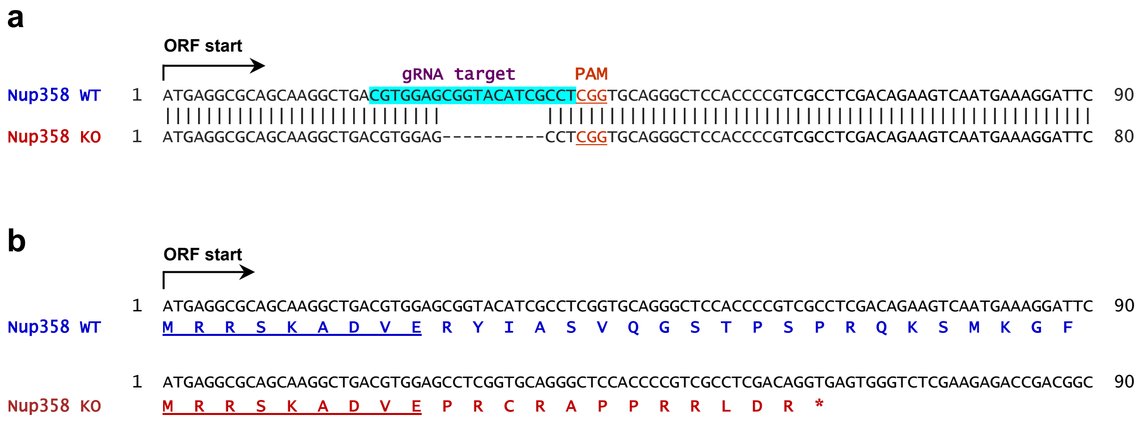


**Supplementary Fig. 3 | Details of Nup358 KO HeLa cell line. a**, HeLa cells were transfected with a construct encoding Cas9 protein, and a guide(g) RNA targeted against Nup358 (highlighted in blue in Nup358 WT sequence). PAM sequence in Nup358 is also labelled. The Nup358 knockout (KO) line that was used in this study was heterozygous. It had one wild type allele and the other allele had deletions (indicated as dashed lines in Nup358 KO sequence). **b**, The KO cell line used in this study has a 10-nucleotide deletion in one of the alleles, thus rendering it incapable of producing full length Nup358, but predicted to produce a 21-amino acid peptide as indicated in Nup358 KO sequence. Common sequence present in the WT and peptide derived from the KO allele are underlined. * indicates stop codon. ORF; open reading frame.

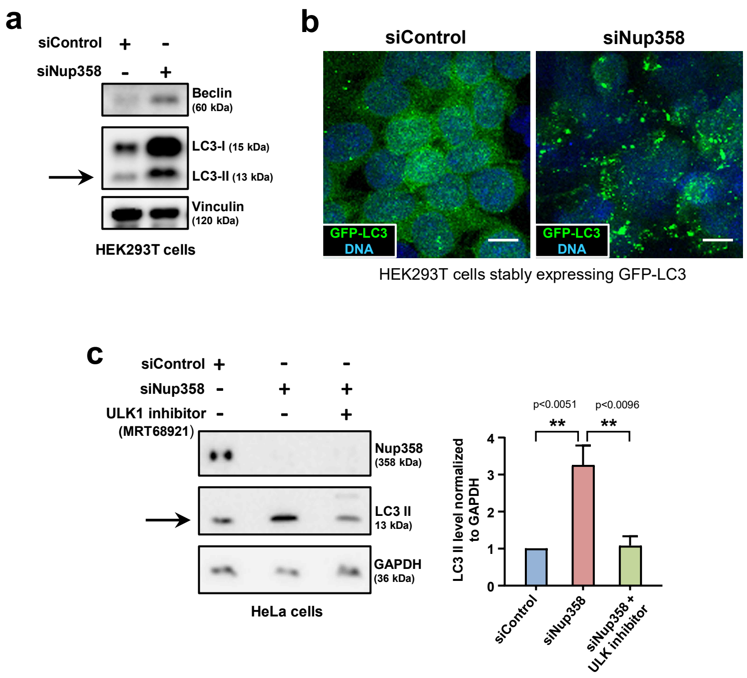


**Supplementary Fig. 4 | Nup358 deficiency induces autophagy. a**, Depletion of Nup358 induces autophagy. HEK293T cells were treated with siControl or siNup358 and monitored for autophagy by analysing the levels of indicated proteins by western blotting. Extent of Nup358 depletion and levels of different autophagy markers were examined using specific antibodies. Vinculin was used as loading control. Arrow indicates LC3-II. **b**, HEK293T cells stably expressing GFP-LC3 were transfected with siControl or siNup358 as indicated and LC3 puncta formation was visualized by fluorescence microscopy. Scale bar, 10 µm. **c**, Induction of autophagy in Nup358-deficient cells depends on ULK1/2. HeLa WT and Nup358 KO cells were treated with vehicle control or ULK1/2 inhibitor (MRT68921, 20 µM) for 1 h. Left: cells were then lysed and analysed for LC-II levels (indicated by arrow) by western blotting to monitor autophagy. Right: LC-II levels were quantitated in the indicated samples (*n* = 3 independent experiments). Data are mean±SEM, unpaired Student's *t* test. ** $P \leq 0.01$.

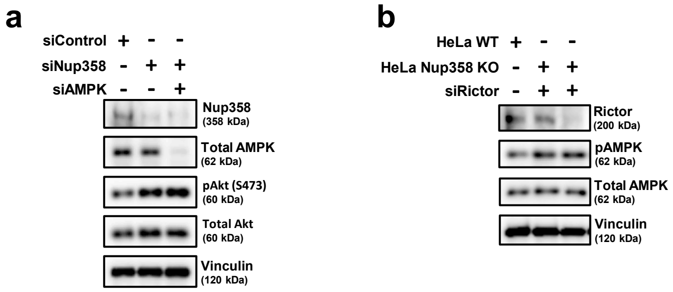


**Supplementary Fig. 5 | Nup358 appears to restrict activation of mTORC2/Akt and AMPK through independent mechanisms. a**, mTORC2/Akt activation in Nup358-deficient cells occur independent of AMPK. HeLa cells were treated with specfic siRNAs as shown and analysed for the activation status of mTORC2/Akt indicated by phosphorylation of Akt at S473 by western blotting using indicated antibodies. α-tubulin was used as loading control. **b**, AMPK activation upon Nup358 depletion is independent of mTORC2/Akt signalling. HeLa WT and Nup358 KO cells were transfected with control (siControl) or Rictor specific (siRictor) siRNA and analysed for the activation status of AMPK by monitoring its phosphorylation at T172 by western blotting. Vinculin was used as loading control.
